# Diversity and composition of microbial communities in shrimp ponds sediments: A metagenomic approach

**DOI:** 10.1101/2025.08.29.672896

**Authors:** Telmo A. Escobar-Troya, J. José Palma-Pozo, José Flores, Katiuska López, Samir Zambrano, Camilo Mestanza

## Abstract

The cultivation of *Litopenaeus vannamei* is of great economic importance to Ecuador, yet it faces significant challenges due to microbial diseases. This study analyzes the microbial metagenome in white shrimp culture ponds sediments, assessing the diversity and communities of both pathogenic and non-pathogenic bacteria. Sediment samples were collected from ponds in three provinces with varying salinity levels: province of Guayas (6.28 g/L), El Oro (21 g/L), and Santa Elena (48 g/L). Through sequencing the V3-V4 region of the 16S rRNA gene, diverse microbial communities were identified, significantly influenced by salinity. Results revealed that microbial communities vary among locations, highlighting the presence of pathogenic bacteria such as Proteobacteria and beneficial ones like Firmicutes. Shannon and Simpson indices indicated high diversity and evenness in Guayas, El Oro, and Santa Elena (Shannon: 5.547, 5.810, and 4.326; Simpson: 0.9953, 0.9957, and 0.982, respectively). This analysis provides crucial information for improving management and sustainability of shrimp aquaculture, offering insights into the complex microbial ecosystems in culture environments and their potential impact on shrimp health and production.

## INTRODUCTION

The cultivation of white shrimp (*Litopenaeus vannamei*) represents a significant economic activity in Ecuador, yet its productivity is challenged by several factors, diseases caused by microorganisms (archaea, bacteria, protozoa) and viruses (HUANG L et al.,2011). These pathogens dramatically affect the species’ growth process, with reports estimating that costs associated with controlling losses in the shrimp industry due to microbiome-related pathologies range between 10,000 and 15,000 USD/ha (TERREROS PONCE, F. A, 2025).

*L. vannamei* cultivation methods vary from extensive, utilizing large ponds areas with low density, to intensive systems characterized by high population densities in smaller ponds with rigorous control of environmental factors and feeding (PEREIRA SANTOS et al., 2020). However, these practices can lead to dysbiosis - changes in the composition and abundance of microbial communities (PRIMAVERA JH, 2006). Factors such as salinity levels, high population densities, overfeeding, and chemical use can significantly alter microbial structure, leading to imbalances affecting soil quality and organism health (CORNEJO-GRANADOS F et al., 2018).

Water salinity plays a crucial role in the composition and dynamics of microbiome communities. Salinity levels can increase competition between autotrophic and heterotrophic bacteria due to nutrient availability and pH changes. Non-pathogenic soil bacteria like *Bacillus* spp., *Lactobacillus* spp., *Nitrosomonas* spp., and *Nitrobacter* spp. activate various metabolic mechanisms, including organic matter degradation, antimicrobial substance production, and nitrogen cycle balancing (WEIXING & JUNYUAN, 2020; FEITO J., et al 2022). Conversely, pathogenic bacteria species, notably *Vibrios* spp. and *Aeromonas* spp., thrive in both low and high salinities, causing severe diseases like acute hepatopancreatic necrosis syndrome (DHANUSH, G., 2025).

Metagenomic studies of *L. vannamei* digestive tract have revealed both positive and negative impacts of microbial diversity on shrimp growth and metabolic performance. Some bacteria, such as *Shimia marina* and *Vibrio* sp., are associated with slower growth, while others like *Candidatus bacilloplasma, Tamlana agarivorans*, and *Donghicola tyrosinivorans* are linked to faster growth and improved metabolic yield (JIA-JUN et al., 2019; LANFEN FAN & QING X. Li, 2019).

Metagenomic analysis allows for the creation of robust molecular patterns, facilitating the specific characterization of genomic profiles in situ. This tool can significantly contribute to decision-making and lead to the development of technical management strategies in the field (SCHLOSS PD & HANDELSMAN J, 2003).

This study analyzes the metagenomic gradient of bacterial 16S rRNA from sediment samples in *Litopenaeus vannamei* production ponds. The primary aim is to determine bacterial diversity by establishing the main pathogenic and non-pathogenic microbial communities (TARNECKI AM et al., 2017). This research will provide crucial insights into the complex microbial ecosystems in shrimp aquaculture environments, potentially improving management practices and sustainability in this economically vital industry.

## MATERIALS AND METHODS

### Description of the study area

This study was conducted in three provincial locations in Ecuador (Figure 2), focusing on shrimp farming ponds of *Litopenaeus vannamei*. The sampled sites included Puerto Roma (Guayas - 2°30’52.6"S 79°53’06.6"W), Colonche (Santa Elena - 2°00’23.2"S 80°43’11.4"W), and El Retiro (El Oro - 3°24’26.9"S 79°58’50.7"W). These locations were selected due to their salinity gradient, environmental conditions, and farming practices (ZENG et al., 2020). Shrimp ponds in each province were chosen and classified according to salinity levels: low (≤ 15 g/L), medium (16-30 g/L), and high (≥ 30 g/L).

### Determination of water and sediment quality

Water quality parameters such as pH, were measured *in situ* using strips (Microessential Laboratory), salinity using a portable optical refractometer (S-100), and temperature using a portable digital thermometer (Vetco Supply, USA). A sediment corer was used for sample collection at each site. Three random samples were taken from the selected ponds, using a corer approximately 15 cm in diameter and two meters in length. The corer was vertically submerged into the water to a depth of approximately two meters, allowing the collection of soil samples from an approximate depth of 10 cm. A total of 10 grams of soil were extracted per sample and placed into sterile 15 mL Falcon tubes. The samples were labeled according to the sampling location with the following code: sampling 1 Province. del Guayas - Puerto Roma «PRP1, PRR2, PRP3, sampling 2 Province. El Oro - El Retiro «ERP1, ERP2, ERP3, and sampling 3 Province Santa Elena - Colonche (COP1, COP2, COP3). The tubes were hermetically sealed and stored in a climate-controlled cooler at approximately 4°C before being transported to the Biotechnology Laboratory at the University of Guayaquil, Faculty of Natural Sciences.

### Extraction, amplification and sequencing of bacterial DNA from sediment

The samples were preserved at -10°C and were divided into two groups for DNA extraction. The ZymoBIOMICS DNA (Zymo Research, USA) extraction kit was used to extract nucleic acids according to the manufacturer protocol.

For the identification and characterization of bacterial diversity. for this, were used the sequences reported for the Illumina MiSeq System. 16S Amplicon PCR, labeled with barcodes containing random oligos of 6 bases. Forward Primer 341: 5’CCTACGGGNGGCWGCAG3’ and Reverse Primer 805 5’GACTACHVGGGTATCTAATCC3’, synthesizing a single amplicon of ∼460 bp. As directed by the parent laboratory, the PCR test was carried out in a thermocycler following the thermocycling programmed: 95°C for 3 minutes for initial denaturation, followed by 25 cycles of 95°C for 30 seconds, 55°C for 30 seconds and 72°C for 30 seconds, with a final extension at 72°C for 5 minutes (ILLUMINA, 2013). After amplification, PCR products were purified using magnetic beads to remove residual primers and other contaminants. DNA quantity and quality were analyzed by spectrophotometry (ND-1000-NanoDrop Technologies Wilmington, USA).

For the preparation of gene libraries and arrays, unique sequencing indices were added to each end of the amplicons by a second round of PCR. Once indexed, the amplicons were subjected to a second wash with magnetic beads to ensure cleanliness of the final library. Sequencing was performed under the Illumina MiSeq platform (Illumina, California, USA.) The raw data generated from the MiSeq platform were paired-end readings.

### Bioinformatics and statistical analysis

The sequences sent by the commercial company of all sediment samples were processed, considering the sequences readings with 2% error and not aligned, which were not considered for the analysis. The files transformed to Fastq that were placed in the Drive. using Google Colab Pro. On the other hand, to measure the alpha diversity of the bacterial communities, the Shannon and Simpson index were included, using the QIIME2 platform. The Bray-Curtis distance was used to evaluate the differences in species complexity of the samples.

For molecular taxonomic classification from the identified ASVs, the trained comparative database for the Neive Bayes classifier of “SILVA” was used with the default confidence threshold at ≥ 0.5. As for the OTU abundance information, the data was normalized by using a sequence number standard corresponding to the sample with the fewest sequences. A principal coordinates analysis (PCoA), was performed to demonstrate the clustering of samples using the “vegan” and “phyloseq” packages. To evaluate salinity conditions, these were exempted from statistical indicators. The NCBI platform was used to analyze the relationship between pathogenic and non-pathogenic microbial communities present in the metagenomic sequences examined.

## RESULTS AND DISCUSSION

The culture of *Litopennaeus vannamei* is a significant resource in Ecuador; however, little attention has been given to the molecular identification of the accompanying microbiomes in its production ecosystem. The interpretation of the interaction of microorganisms with the ecosystem and the feeding behavior of the animals is a potential tool for the efficient management of the culture of this species. In our study, we compared the presence of pathogenic and non-pathogenic microbial groups in three shrimp farms with variable salinity gradients and high commercial production. Environmental factors in samples in the provinces of Guayas were «6.28 g/L-∇=0.29; pH 9.08; 34.8 °C », Santa Elena «21g/L-∇=0,61; pH 6.7; 34 °C» and El Oro «48 g/L-∇=1; pH 7.2; 34.2 ». These differences, especially that of salinity, made it possible to evaluate microbial differentiation in *L. vannamei* culture (COLETTE et al., 2023).

### Microbial composition and diversity

Based on analysis of the demultiplexed sequences and quality and chimera filtering of the raw data, a total average of 155336 forward and reverse reads was produced. The quality indices suggest that each sequenced sample presented acceptable conditions at the molecular level for the metagenomic study.

According to alpha diversity analysis, it showed significant differences in diversity and richness indices. Good coverage approached 100% in all samples, indicating that most of the species present were detected, ensuring an adequate sequencing depth and a complete representation of the microbial communities. Shannon and Simpson indices showed that Guayas and El Oro have higher diversity and equity (Shannon: 5.547 y 5.810; Simpson: 0.9953 y 0.9957). The ACE and Chao1 richness indices highlighted El Oro as the most diverse community (ACE: 522,156; Chao1: 522) in addition, the high presence of Cyanobacteria is common in aquatic environments with moderate levels of salinity, where they can fix nitrogen and produce toxins (ZHANG et al., 2019). These data suggest that the differences may be influenced by environmental factors and aquaculture management practices (HOU et al., 2017).

### Comparative analysis of frequent taxonomic groups

In the samples from the Guayas shrimp farm, they showed a high abundance of the Proteobacteria group, a diverse phylum that includes both pathogenic and non-pathogenic bacteria, such as those of the genus *Vibrio*, known to cause disease in shrimp (ZHOU et al., 2024). Bacteroidetes, which aid in the degradation of organic matter, and Firmicutes, including Bacillus, which may act as probiotics, were also found, while the samples from El Oro were dominated by Bacteroidetes, Cyanobacteria and Proteobacteria, suggesting a community specialized in the degradation of organic compounds (LAN ANH et al., 2020), although the presence of Proteobacteria could indicate potential pathogens (ZHANG et al., 2019). Samples from the Santa Elena shrimp farm showed a higher relative diversity, with a balanced distribution of Proteobacteria, Bacteroidetes, Actinobacteria and Firmicutes, suggesting an improvement in the quality of the pool soil and a healthy environment for shrimp (QIAN et al., 2022). Water salinity significantly influences microbial composition, with Proteobacteria predominating (KRISTIN et al., 2019). El Oro with Bacteroidetes and Cyanobacteria, and Guayas with higher microbial diversity, indicate that variability in salinity affects the microbial community structure in these culture pools (Figure 1).

**Figure 1.**
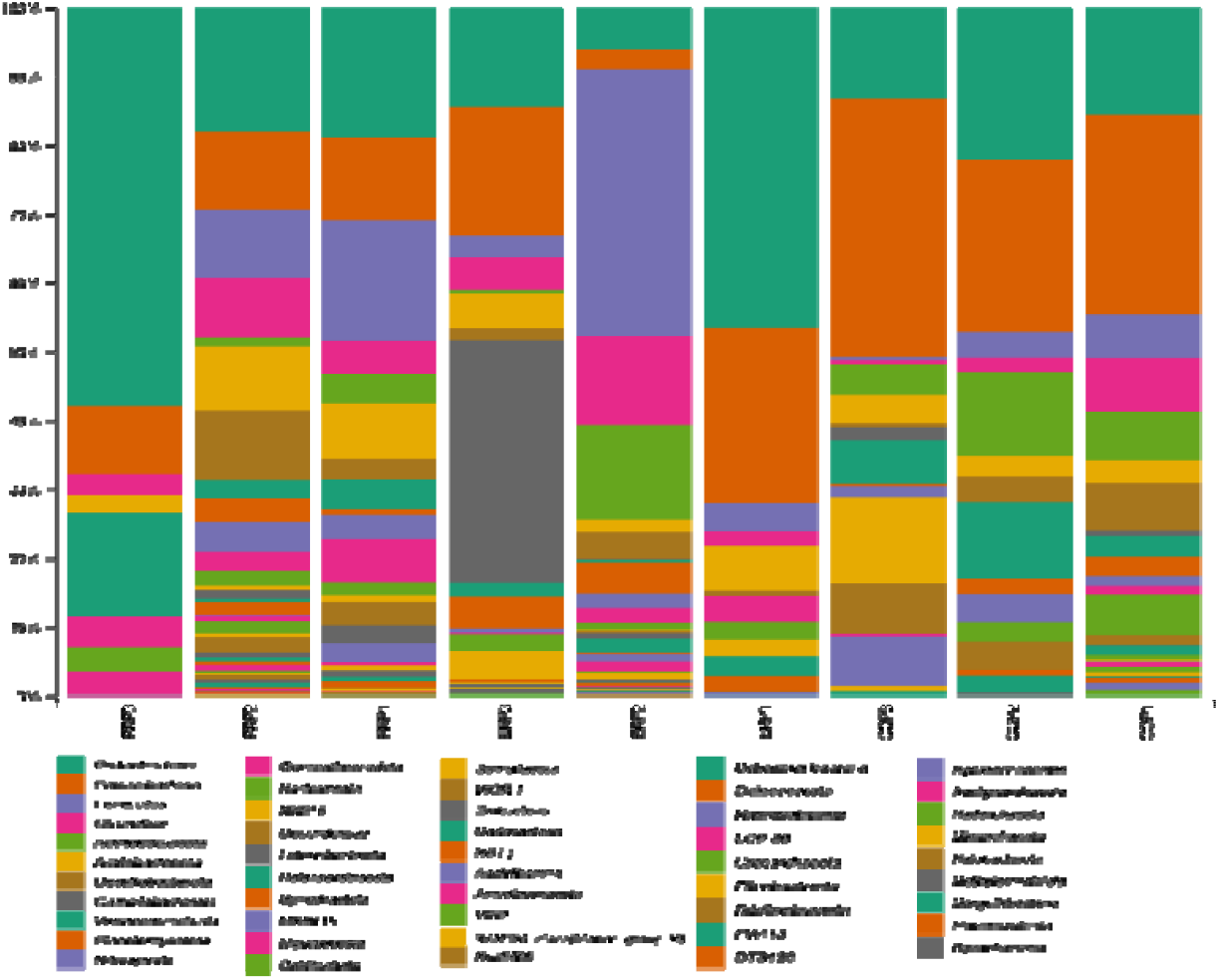
Abundance in the sediment samples from Guayas (PRP1, PRR2, PRP3), El Oro (ERP1, ERP2, ERP3) y Santa Elena (COP1, COP2, COP3).

### Evidence of a shared bacterial community associated with sampling sites

A total of 1,568 unique microbial genera were identified in the three provinces: El Oro 704 (44.8%), Guayas 500 (31.8%), and Santa Elena 364 (23.2%). The diagram showed three shared OTUs that were associated with all sites examined (Figure 2a). In context, significant variations in the composition of microbial communities are demonstrated in all samples analyzed. The relationship between specific samples and taxonomic groups of bacteria is distributed according to colors ranging from yellow (highest abundance) to black (lowest abundance), indicating the relative presence of different microbial groups in each sample. The upper dendrogram of the hierarchical clustering of the heat map indicates similar microbial compositions in the samples from El Oro and Guayas, in addition to showing a greater diversity and abundance of bacteria, among those of relevance are Halobacillus and Bacillus type, the presence of them suggests positive roles in animal health (ERCHAO LI et al., 2018; PINOARGOTE et al., 2018). In addition, the presence of Pseudomonas is considered as indicators of effective recovery and biodegradation of complex organic compounds (FARIDUDDIN et al., 2018), including contaminants (COLETTE et al., 2023). Also, the genera Candidatus and Woesebacteria indicate their possible positive ecological importance in these environments (KRISTIN et al., 2019). However, the presence of Escherichia and Shigella in sediments may be an indicator of possible fecal contamination (PRIYADARSANI DAS et al., 2018). In contrast, the El Oro samples clustered differently, indicating significant differences in their microbial communities compared to the other samples (Figure 2b). *In situ* analysis of this type of sample, allows determining the microbial community, which when balanced in the sediment can improve feeding efficiency, reduce mortality and increase biomass production.

**Figure 2.**
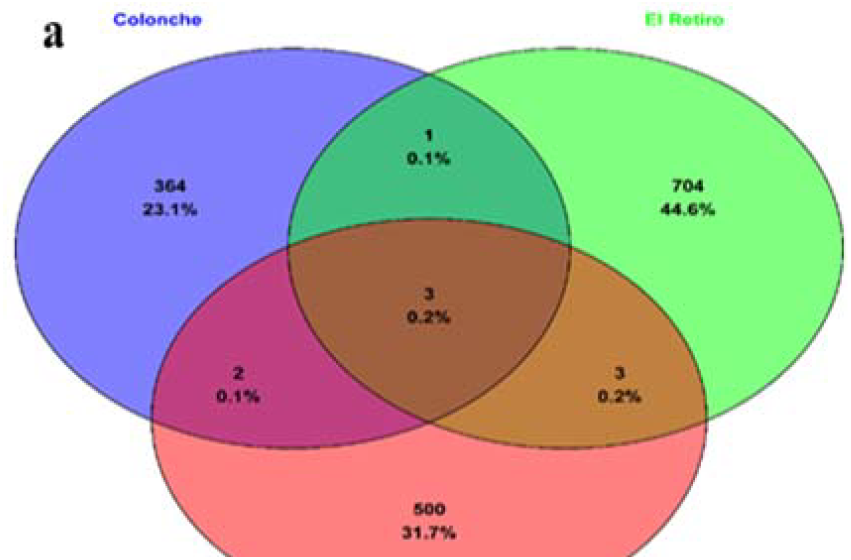
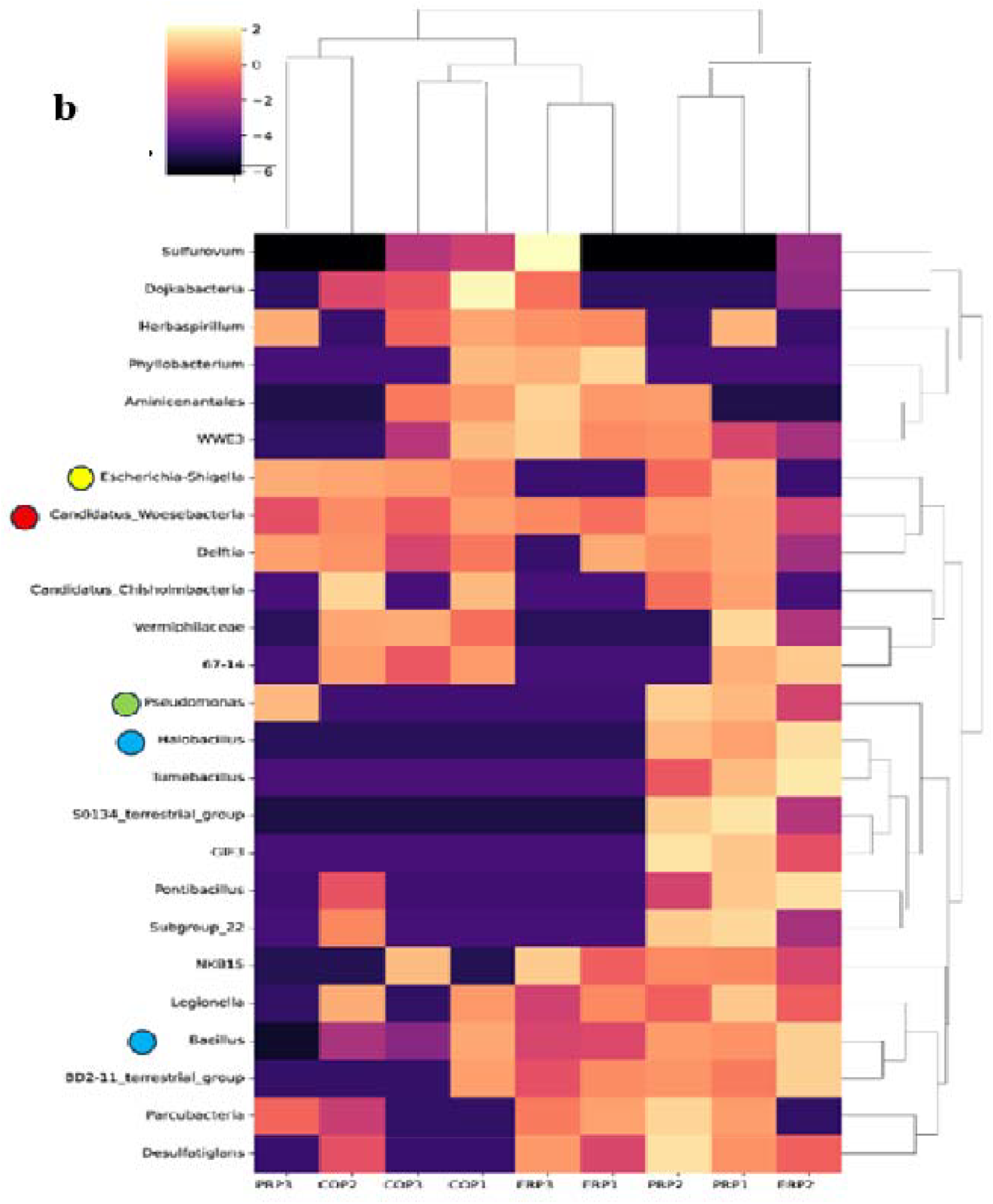
The Venn diagram shows the number of OTUs that were unique or shared at the sampling sites (**a**). The heat map represents the relative abundance of unique or shared bacterial taxa at the OTUS level in sediment samples from the sampling sites. (**b**).

### Beta diversity analysis

Based on the Bray-Curtis index, it showed significant differences in microbial community composition among the different shrimp pond sediment samples, which were strongly influenced by water salinity (LI et al., 2017). Samples PRP1, PRP2 and PRP3 showed dissimilarity ≥ 0.987 and 1, with PRP2 having the highest dissimilarity 1, suggesting a quite different microbial composition. Meanwhile, ERP1 and PRP3 showed a < 0.926, indicating greater similarity in their microbial communities. However, ERP2 showed a high dissimilarity (1) with several samples, as did PRP2. As for the Santa Elena code samples, COP1 and COP2 showed >0.971, although COP2 showed maximum dissimilarity (1) with PRP2, ERP2 and COP3. Non-metric multidimensional scaling (NMDS) analysis based on the Bray-Curtis index reveals that a salinity of 21 g/L (blue) shows greater variability and does not form a cohesive group (Figure 3). These data suggest robust variations in microbial composition and structure in the three locations, i.e., due to differences in salinity (MACEDO DB et al., 2024).

**Figure 3.**
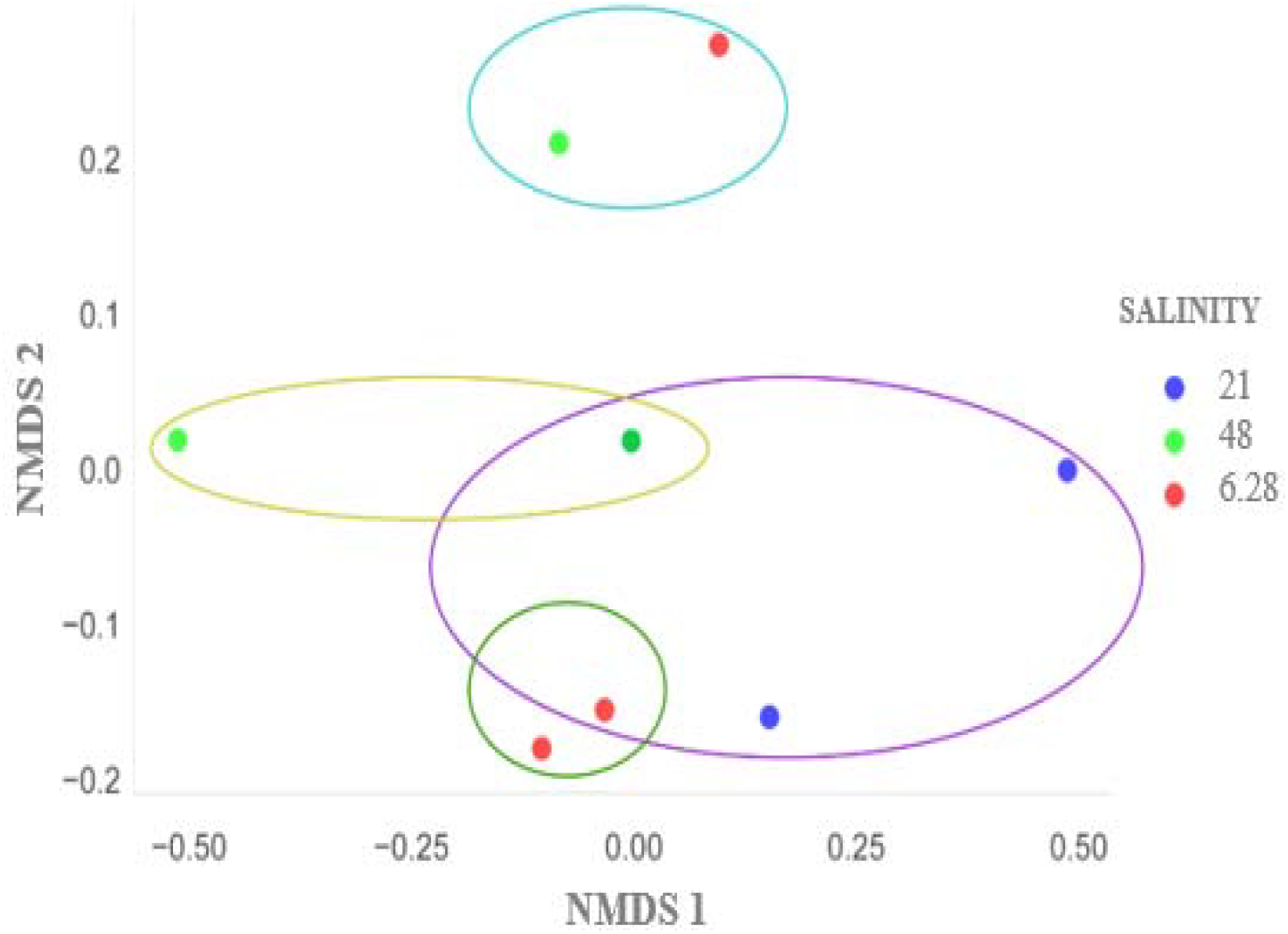
Non-metric multidimensional scaling (NMDS) analysis based on the Bray-Curtis index mediated by salinity and its impact on microbial composition in sediments.

### Pathogenic bacterial analysis

The taxa and percentage of metagenome reads found are shown in table 1. We determined the significance of coefficients (P <0.05). No significant statistical differences P=0.3954, between Guayas (71633), El Oro (55801) and Santa Elena (14134). In context, the Pseudomonas group represented 12.02% in the Guayas sample, the presence of this group is an indicator of vital importance in production pools (CAI D et al., 2021), since these genera contain the toxA gene, which encodes exotoxin A and its action of inhibiting protein synthesis in host cells causes high mortality rates in the pools (GERALDINE DURANGO et al., 2024). Among the other groups of interest found, is the genus Acinetobacter with 2.40%, these bacteria possess the aprA and ompA genes (SCRIBANO D et al., 2024), the former encodes the alkaline protease which contributes to the degradation of shrimp tissues and the latter which favors adhesion and evasion of the immune system of the organism (XUAN DONG et al., 2019).

**Table 1.** Composition and distribution of the dominant pathogenic bacterial microbiota in sediment from shrimp farms in the provinces of Guayas, El Oro, and Santa Elena.

| Shrimp farm |  | Taxonomy | % readings |
| --- | --- | --- | --- |
| Guayas | Proteobacteria | Acinetobacter | 2.40 |
|  |  | Pseudomonas | 12.02 |
|  | Actinobacteriota | Mycobacteriaceae | 0.31 |
|  | Gemmatimonadota | BD2-11 | 1.25 |
|  |  | Gemmatimonadetes | 0.37 |
| El Oro | Proteobacteria | Acinetobacter | 0.03 |
|  |  | Pseudomonas | 1.39 |
|  | Actinobacteriota | Mycobacteriaceae | 0.69 |
|  | Gemmatimonadota | BD2-11 | 5.20 |
|  |  | Gemmatimonadetes | 0.01 |
| Santa Elena | Proteobacteria | Acinetobacter | 0.02 |
|  |  | Pseudomonas | 0.03 |
|  | Actinobacteriota | Mycobacteriaceae | 0.01 |
|  | Gemmatimonadota | BD2-11 | 1.22 |
|  |  | Gemmatimonadetes | 0.02 |

The presence of the Mycobacteriaceae family is notable with 0.31% of the readings in Guayas (0.69%), El Oro and Santa Elena (0.01%). It is a group of interest in the pathology of crustaceans, because this microbial group contains the hsp65 gene (RINGUET H et al., 1999), which encodes a heat shock protein that enables bacteria to survive in adverse conditions, and its presence is usually associated with chronic melanized lesions in the cuticle and internal organs, without causing great mortalities (JACOBS, J et al., 2009). Transmission is horizontal, affecting adult shrimp (MERCE BERGA et al., 2017).

The phylum Gemmatimonadota identified in the samples showed significant percentages with readings of the genus BD2-11, which were for El Oro (5.20%), Santa Elena (1.22%) and Guayas (1.25%). This group is little known for its pathogenicity, however, the high presence of BD2-11 suggests effective adaptation and cell amplification. Regular monitoring in the shrimp culture environment because its uncontrolled amplification could dramatically influence the shrimp microbiota and its immune response LAN ANH et al. (2020). The common denominator of these groups analyzed is that, according to the works reviewed here, all of them can directly influence the immune response of shrimp (XIONG J et al., 2017).

### Nonpathogenic bacteria analysis

In this context, the percentage of readings are shown in table 2. Bacteria of the phylum Firmicutes, the Bacilli class was abundant in all samples, with readings of 20.72% in Guayas, El Oro (25.58%), and Santa Elena with 7.24%. Within this class, these are known for their ability to break down complex organic matter into simpler compounds through fermentation CHITHIRA et al. (2021), effectively utilizing enzymatic synthesis such as hydrolases and dehydrogenases (JIA JUN et al., 2019). In the soil of shrimp ponds, the decomposition of organic matter is particularly important for its mineralization ZIFANG CHI et al. (2021), which enhances the carbon cycle and improves the availability of nutrients (QIAN, L. (2022). Among other potential enzymatic activities of this class Bacilli possesses the ability to fix atmospheric nitrogen HOU et al. (2017), converting it into ammonia through nitrogenase activity LEI WANG et al. (2022).

**Table 2.**
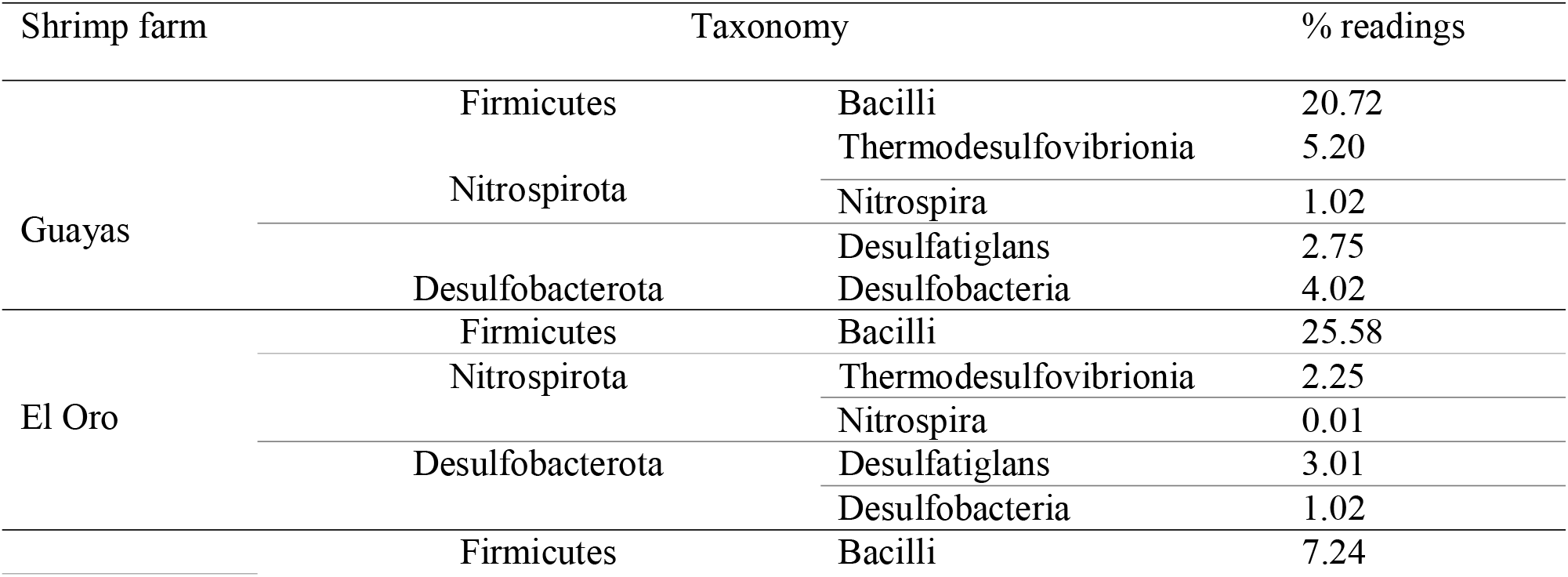
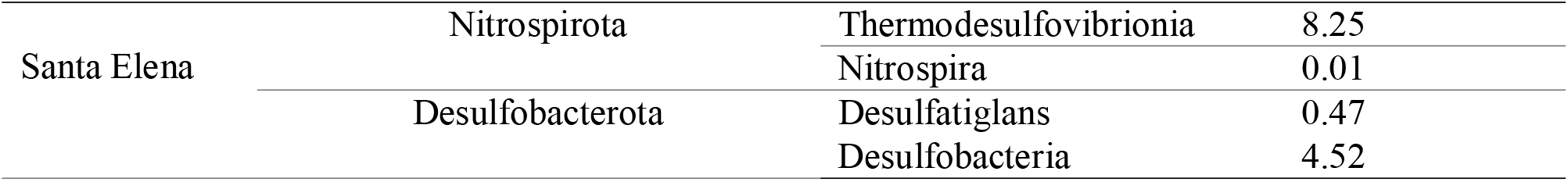
Composition and distribution of the dominant non-pathogenic bacterial microbiota in sediment from shrimp farms in the provinces of Guayas, El Oro, and Santa Elena.

The findings of the Nitrospirota phylum, Nitrospira and Thermodesulfovibrionia classes, are positive indicators in the sediment of shrimp ponds (UMEZAWA K et al., 2021). Based on the examination, Nitrospira presented 1.02% in Guayas, while in smaller amounts in El Oro and Santa Elena. Microbial activity in the nitrogen cycle is critical. Most of the species in this group are involved in nitrification through the enzymatic activity of nitrite oxidoreductase, which causes the oxidation of NO^2^ to NO^3^ (XURI DONG et al., 2023).

Thermodesulfovibrionia and Desulfobacterota participate in the reduction of SO^4^_2_ to H_2_S (LAN ANH et al., (2020), this being a key process in the sulfur cycle (CHITHIRA et al., 2021). This process is carried out by the action of enzymes such as sulfate reductase, which facilitates the reduction of sulfate to sulfide (COLETTE et al., 2023), integrating sulfur into organic compounds and improving the availability of nutrients in the soil (XIAOHONG et al., 2021).

## CONCLUSION

The microbial composition varies significantly between Santa Elena, El Oro, and Guayas, harboring a unique combination of pathogenic and non-pathogenic bacteria. Salinity is a determining factor that affects the structure and diversity of bacterial communities. In addition, the presence of specific microbial genera is evident at each site, suggesting a close relationship between environmental conditions and microbial ecology.

In Santa Elena, high salinity favors the predominance of Proteobacteria, while in El Oro, Bacteroidetes and *Cyanobacteria* spp. are observed, known both for their nitrogen fixation capacity and their potential for toxin production. In Guayas, with the lowest salinity, greater overall microbial diversity was found. The identification of specific genera, such as *Pseudomonas* sp., *Halobacillus* spp., and *Bacillus* spp. in Guayas and El Oro suggests crucial roles in bioremediation and water quality improvement, indicating potential ecological and practical importance in the management of these aquaculture systems.

## ACKNOWLEDGEMENTS (required)

The authors would like to thank Opúsculo del Mar S.A., the shrimp farming communities of Puerto Roma and El Retiro, to the manager of the CDI-Center for Diagnosis and Biotechnological Research, Francisco Peralta, MSc., for their kind collaboration. The study was funded entirely by the authors of this study.

## DECLARATION OF CONFLICT OF INTEREST (required)

The authors declare no conflict of interest. The founding sponsors had no role in the design of the study; in the collection, analyses, or interpretation of data; in the writing of the manuscript, and in the decision to publish the results.

### Other options

*We have no conflict of interest to declare*.

## AUTHORS’ CONTRIBUTIONS (required)

All authors contributed equally to the conception and writing of the manuscript. All authors critically revised the manuscript and approved of the final version.

## BIOETHICS AND BIOSSECURITY COMMITTEE APPROVAL

**(Required when animals and genetically modified organisms are involved)**

